# grepq: A Rust application that quickly filters FASTQ files by matching sequences to a set of regular expressions

**DOI:** 10.1101/2025.01.09.632104

**Authors:** Nicholas D. Crosbie

## Abstract

Regular expressions (regex) (Kleene 1951) have been an important tool for finding patterns in biological codes for decades (Hodgman 2000 and citations therein), and unlike fuzzy-finding approaches, do not result in approximate matches. The performance of regular expressions can be slow, however, especially when searching for matching patterns in large files. *grepq* is a Rust application that quickly filters FASTQ files by matching sequences to a set of regular expressions. *grepq* is designed with a focus on performance and scalability, is easy to install and easy to use, enabling users to quickly filter large FASTQ files, to enumerate named and unnamed variants, to update the order in which patterns are matched against sequences through in-built *tune* and *summarise* commands, and optionally, to output a SQLite file for further sequence analysis. *grepq* is open-source and available on *GitHub* and *Crates*.*io*.

## Statement of need

The ability to quickly filter FASTQ files by matching sequences to a set of regular expressions is an important task in bioinformatics, especially when working with large datasets. The importance and challenge of this task will only grow as sequencing technologies continue to advance and produce ever larger datasets (Katz et al. 2022). The uses cases of *grepq* are diverse, and include preprocessing of FASTQ files before downstream analysis, quality control of sequencing data, and filtering out unwanted sequences. Where decisions need be made quickly, such as in a clinical settings (Bachurin et al. 2024), biosecurity (Valdivia-Granda 2012), and wastewater-based epidemiology in support of public health measures (Choi et al. 2018; Sims and Kasprzyk-Hordern 2020; Xylogiannopoulos 2021; Merrett et al. 2024), the ability to quickly filter FASTQ files and enumerate named and unnamed variants by matching sequences to a set of regular expressions is attractive as it circumvents the need for more time-consuming bioinformatic workflows.

Regular expressions are a powerful tool for matching sequences, but they can be slow and inefficient when working with large datasets. Furthermore, general purpose tools like *grep* (Free Software Foundation 2023) and *ripgrep* (A. Gallant 2025) are not optimized for the specific task of filtering FASTQ files, and ocassionaly yield false positives as they scan the entire FASTQ record, including the sequence quality field. Tools such *awk* (Aho, Kernighan, and Weinberger 1988) and *gawk* (Free Software Foundation 2024) can be used to filter FASTQ files without yielding false positives, but they are significantly slower than *grepq* and can require the development of more complex scripts to achieve the same result.

## Implementation

*grepq* is implemented in Rust, a systems programming language known for its safety features, which help prevent common programming errors such as null pointer dereferences and buffer overflows. These features make Rust an ideal choice for implementing a tool like *grepq*, which needs to be fast, efficient, and reliable.

Furthermore, *grepq* obtains its performance and reliability, in part, by using the *seq_io* (Schlegel and Seyboldt 2025) and *regex* (Gallant et al. 2025b) libraries. The *seq_io* library is a well-tested library for parsing FASTQ files, designed to be fast and efficient, and which includes a module for parallel processing of FASTQ records through multi-threading. The *regex* library is designed to work with regular expressions and sets of regular expressions, and is known to be one of the fastest regular expression libraries currently available (Gallant et al. 2025a). The *regex* library supports Perl-like regular expressions without look-around or backreferences (documented at https://docs.rs/regex/1.*/regex/#syntax).

Further performance gains were obtained by:

- use of the *RegexSet* struct from the *regex* library to match multiple regular expressions against a sequence in a single pass, rather than matching each regular expression individually (the *RegexSet* is created and compiled once before entering any loop that processes the FASTQ records, avoiding the overhead of recompiling the regular expressions for each record)
- multi-threading to process the records within an input FASTQ file in parallel through use of multiple CPU cores
- use of the *zlib-ng* backend to the *flate2* library to read and write gzip-compressed FASTQ files, which is faster than the default *miniz_oxide* backend
- use of an optimised global memory allocator (the *mimalloc* library (Mutiple, n.d.)) to reduce memory fragmentation and improve memory allocation and deallocation performance
- buffer reuse to reduce the number of memory allocations and deallocations
- use of byte slices to avoid the overhead of converting to and from string types
- in-lining of performance-critical functions
- use of the *write_all* I/O operation that ensures the data is written in one go, rather than writing data in smaller chunks

## Feature set

*grepq* has the following features:

- support for presence and absence (inverted) matching of a set of regular expressions
- IUPAC ambiguity code support (N, R, Y, etc.)
- support for gzip and zstd compression (reading and writing)
- JSON support for pattern file input and *tune* and *summarise* command output, allowing named regular expression sets and named regular expressions (pattern files can also be in plain text)
- the ability to:

- set predicates to filter FASTQ records on the header field (= record ID line) using a regular expression, minimum sequence length, and minimum average quality score (supports Phred+33 and Phred+64)
- output matched sequences to one of four formats (including FASTQ and FASTA)
- tune the pattern file and enumerate named and unnamed variants with the *tune* and *summarise* commands: these commands will output a plain text or JSON file with the patterns sorted by their frequency of occurrence in the input FASTQ file or gzip-compressed FASTQ file (or a userspecified number of total matches). This can be useful for
- optimizing the pattern file for performance, for example by removing patterns that are rarely matched and reordering nucleotides within the variable regions of the patterns to improve matching efficiency
- count and summarise the total number of records and the number of matching records (or records that don’t match in the case of inverted matching) in the input FASTQ file
- bucket matching sequences to separate files named after each regexName with the **–bucket** flag, in any of the four output formats

Other than when the **inverted** command is given, output to a SQLite database is supported with the **writeSQL** option. The SQLite database will contain a table called **fastq_data** with the following fields: the fastq record (header, sequence and quality fields), length of the sequence field (length), percent GC content (GC), percent GC content as an integer (GC_int), number of unique tetranucleotides in the sequence (nTN), percent tetranucleotide frequency within the sequence (TNF), and a JSON array containing the matched regex patterns, the matches and their position(s) in the FASTQ sequence (variants). If the pattern file was given in JSON format and contained a non-null qualityEncoding field, then the average quality score for the sequence field (average_quality) will also be written. The **–num-tetranucleotides** option can be used to limit the number of tetranucleotides written to the TNF field of the fastq_data SQLite table, these being the most or equal most frequent tetranucleotides in the sequence field of the matched FASTQ records. A summary of the invoked query (pattern and data files) is written to a second table called **query**.

Other than when the *tune* or *summarise* command is run, a FASTQ record is deemed to match (and hence provided in the output) when any of the regular expressions in the pattern file match the sequence field of the FASTQ record. Example output of the *tune* command (when given with the **–json-matches** flag) is shown below:

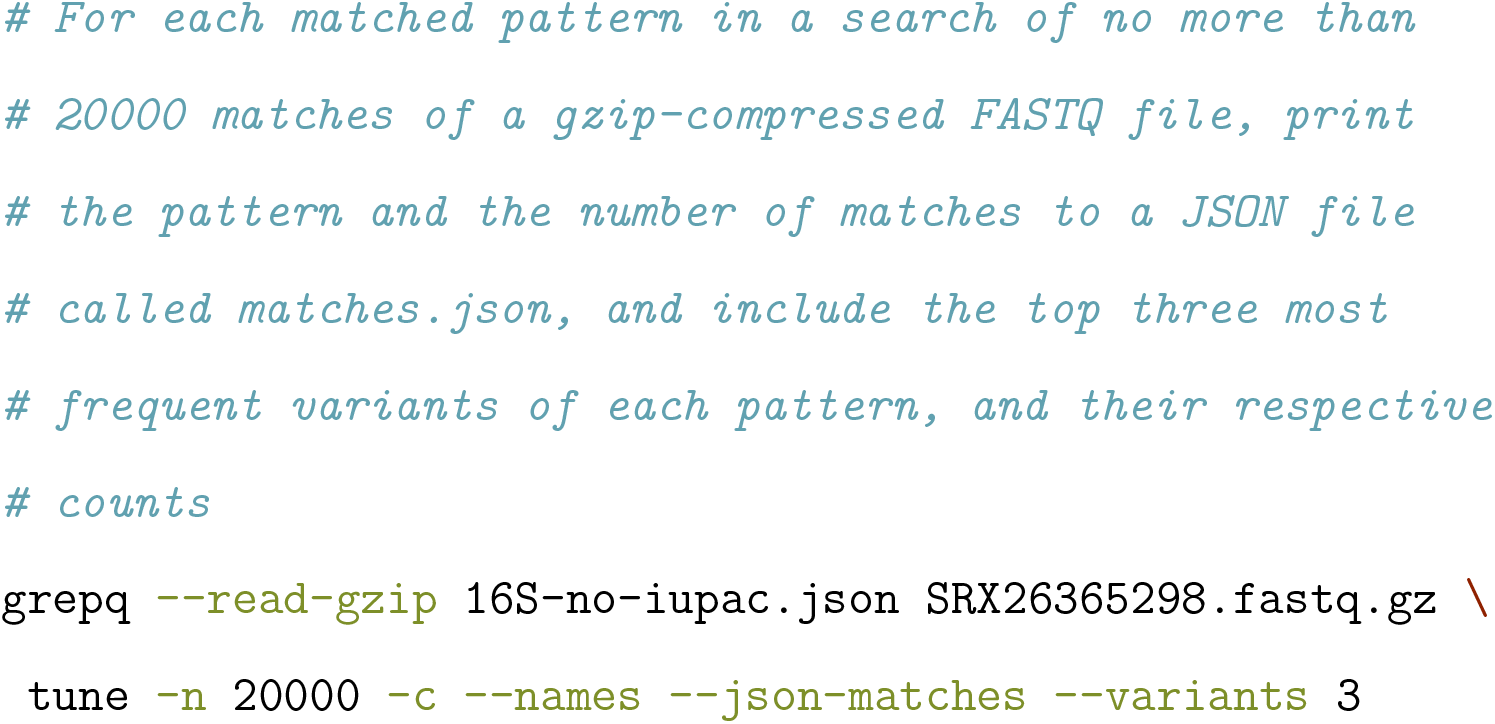

Output (abridged) written to matches.json:

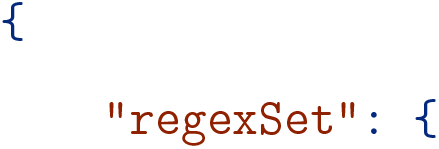
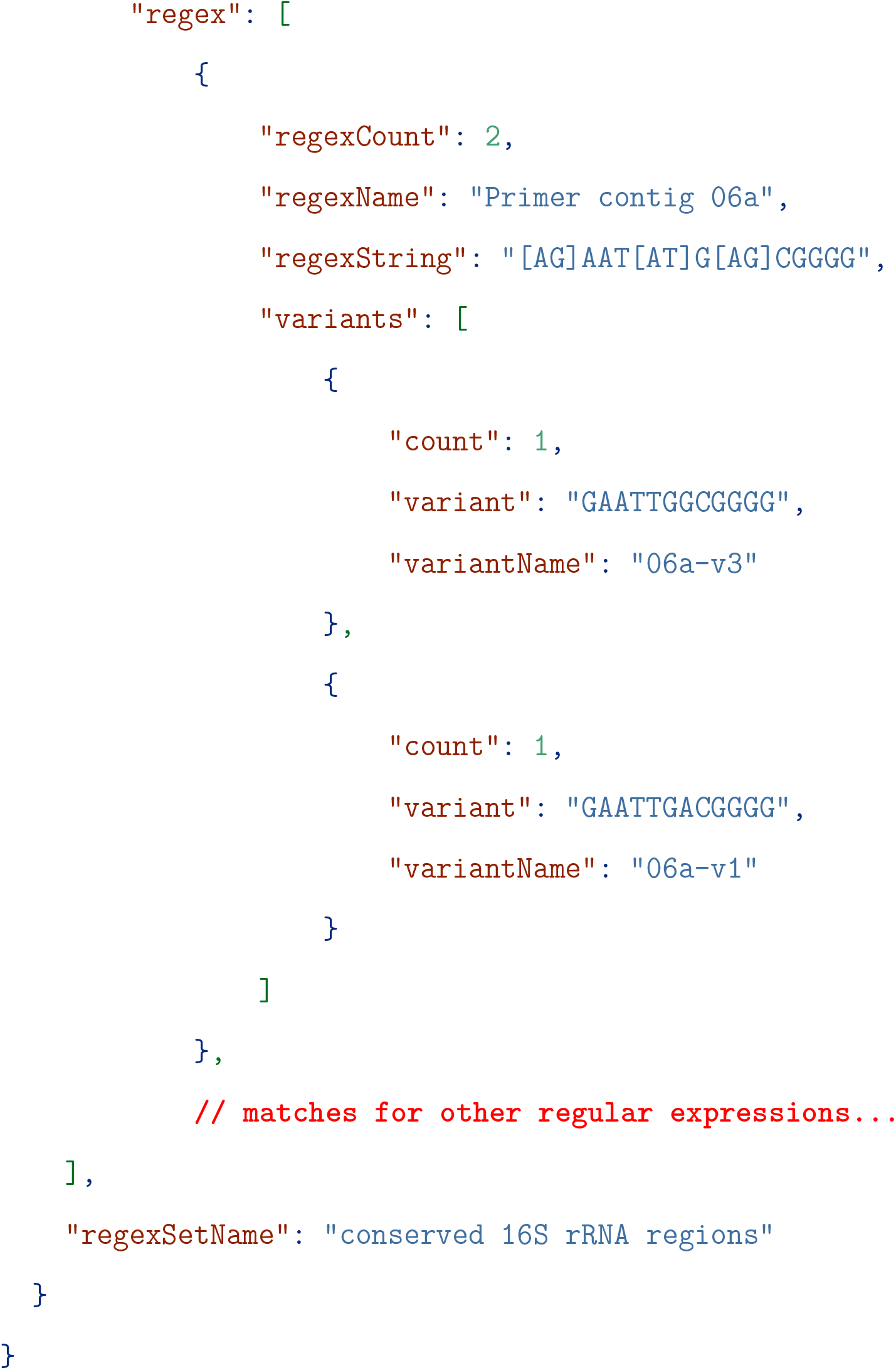

To output all variants of each pattern, use the --all argument, for example:

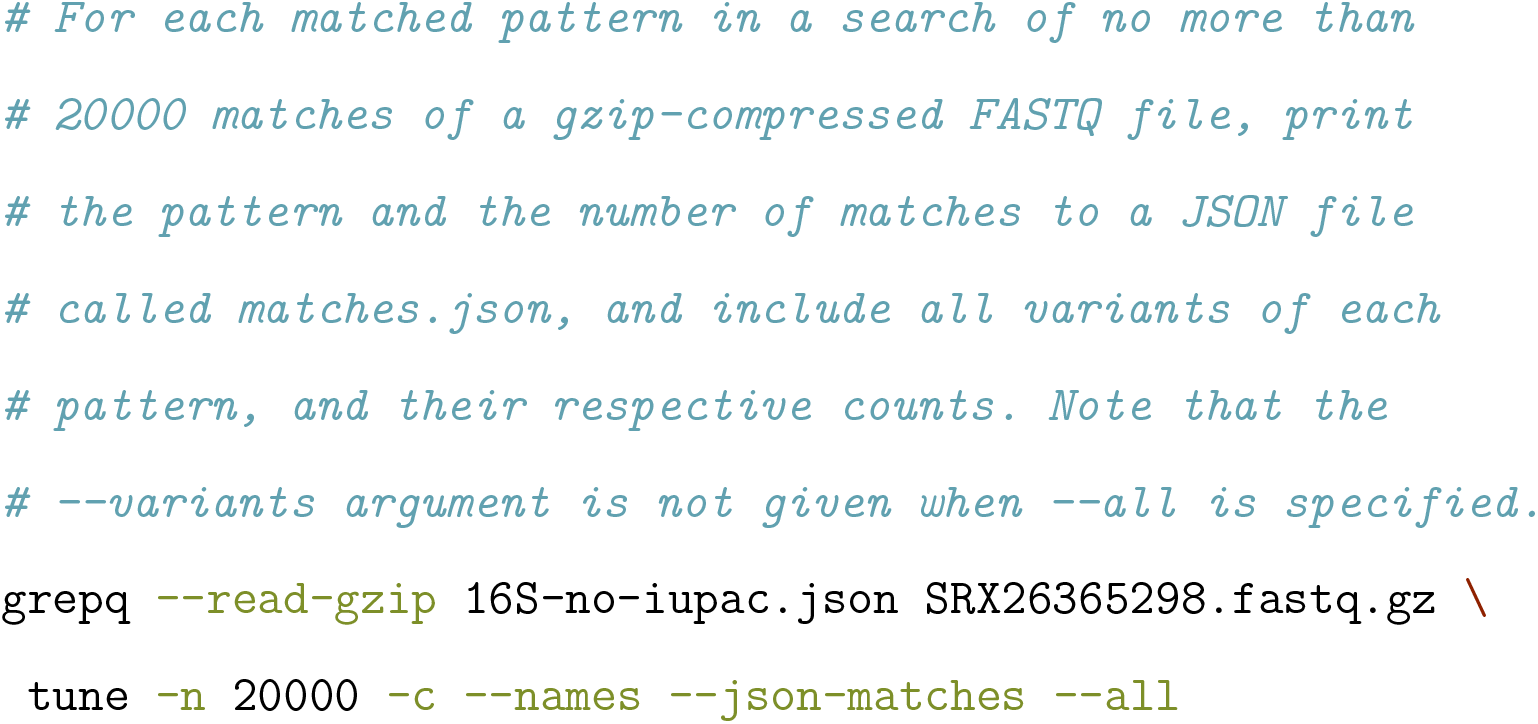

When the count option (**-c**) is given with the *tune* or *summarise* command, *grepq* will count the number of FASTQ records containing a sequence that is matched, for each matching regular expression in the pattern file. If, however, there are multiple occurrences of a given regular expression within a FASTQ record sequence field, *grepq* will count this as one match. To ensure all records are processed, the *summarise* command is used instead of the *tune* command. Further, note that counts produced through independently matching regex patterns to the sequence field of a FASTQ record inherently underestimate the true number of those patterns in the biological sample, since a regex pattern may span two reads (i.e., be truncated at either the beginning or end of a read). To illustrate, a regex pattern representing a 12-mer motif has a 5.5% chance of being truncated for a read length of 400 nucleotides (11/400 + 11/400 = 22/400 = 0.055 or 5.5%), assuming a uniform distribution of motif positions and reads are sampled randomly with respect to motifs (this calculation would need to be adjusted to the extent that motifs are not uniformly distributed and reads are not randomly sampled with respect to motifs).

When the count option (**-c**) is not given as part of the *tune* or *summarise* command, *grepq* provides the total number of matching FASTQ records for the set of regular expressions in the pattern file.

Colorized output for matching regular expressions is not implemented to maximise speed and minimise code complexity, but can be achieved by piping the output to *grep* or *ripgrep* for testing purposes.

## Performance

The performance of *grepq* was compared to that of *fqgrep, seqkit grep, ripgrep, grep, awk*, and *gawk* using the benchmarking tool *hyperfine*. The test conditions and results are shown in **Table 1, Table 2** and **Table 3**.

**Table 1:** Wall times and speedup of various tools for filtering FASTQ records against a set of regular expressions. Test FASTQ file: SRX26365298.fastq (uncompressed) was 874MB in size, and contained 869,034 records.

| tool | wall time (s) |  | speedup |  |  |
| --- | --- | --- | --- | --- | --- |
|  | mean | S.D. | × grep | × ripgrep | × awk |
| *grepq* | 0.192 | 0.010 | 1796.76 | 18.62 | 863.52 |
| *fqgrep* | 0.338 | 0.005 | 1017.61 | 10.55 | 489.07 |
| *ripgrep* | 3.568 | 0.005 | 96.49 | 1.00 | 46.37 |
| *seqkit grep* | 2.885 | 0.011 | 119.33 | 1.24 | 57.35 |
| *grep* | 344.259 | 0.545 | 1.00 | 0.01 | 0.48 |
| *awk* | 165.451 | 1.590 | 2.08 | 0.02 | 1.00 |
| *gawk* | 287.662 | 1.682 | 1.20 | 0.01 | 0.58 |

**Table 2:** Wall times and speedup of various tools for filtering gzip-compressed FASTQ records against a set of regular expressions. Test FASTQ file: SRX26365298.fastq.gz was 266MB in size, and contained 869,034 records.

| tool | wall time (s) |  | speedup |
| --- | --- | --- | --- |
|  | mean | S.D. | × ripgrep |
| *grepq* | 1.703 | 0.002 | 2.10 |
| *fqgrep* | 1.834 | 0.005 | 1.95 |
| *ripgrep* | 3.584 | 0.013 | 1.00 |

**Table 3:** Wall times and speedup of various tools for filtering FASTQ records against a set of regular expressions. Test FASTQ file: SRX22685872.fastq was 104GB in size, and contained 139,700,067 records.

| tool | wall time (s) |  | speedup |
| --- | --- | --- | --- |
|  | mean | S.D. | × ripgrep |
|  | Uncompressed |  |  |
| grepq | 26.972 | 0.244 | 4.41 |
| fqgrep | 50.525 | 0.501 | 2.36 |
| ripgrep | 119.047 | 1.227 | 1.00 |
|  | gzip-compressed |  |  |
| grepq | 149.172 | 1.054 | 0.98 |
| fqgrep | 169.537 | 0.934 | 0.86 |
| ripgrep | 144.333 | 0.243 | 1.00 |

*grepq* v1.4.0, *fqgrep* v.1.02, *ripgrep* v14.1.1, *seqkit grep* v.2.9.0, *grep* 2.6.0-FreeBSD, *awk* v. 20200816, and *gawk* v.5.3.1. *fqgrep* and *seqkit grep* were run with default settings, *ripgrep* was run with **-B 1 -A 2 --colors ‘match:none’ --no-line-number**, and *grep* was run with **-B 1 -A 2 --color=never**. *awk* and *gawk* scripts were also configured to output matching records in FASTQ format. The pattern file contained 30 regular expression representing the 12-mers (and their reverse compliment) from Table 3 of Martinez-Porchas et al. (2017). The wall times, given in seconds, are the mean of 10 runs, and S.D. is the standard deviation of the wall times, also given in seconds.

Test conditions and tool versions as above, but *grepq* was run with the **–read-gzip** option, *fqgrep* with the **-Z** option, and *ripgrep* with the **-z** option. SRX26365298.fastq was gzip-compressed using the *gzip* v.448.0.3 command (Apple Inc. 2019) using default (level 6) settings. The pattern file contained 30 regular expression representing the 12-mers (and their reverse compliment) from Table 3 of Martinez-Porchas et al. (2017). The wall times, given in seconds, are the mean of 10 runs, and S.D. is the standard deviation of the wall times, also given in seconds.

Test conditions and tool versions as described in the footnote to Table 1. Note that when *grepq* was run on the gzip-compressed file, a memory resident time for the *grepq* process of 116M as reported by the *top* command (Apple Inc. 2023c). *fastq-dump* v3.1.1 (Sherry et al. 2012) was used to download SRX22685872 as a gzip compressed file from the NCBI SRA. The pattern file contained 30 regular expression representing the 12-mers (and their reverse compliment) from Table 3 of Martinez-Porchas et al. (2017). The wall times, given in seconds, are the mean of 10 runs, and S.D. is the standard deviation of the wall times, also given in seconds.

## Testing

The output of *grepq* was compared against the output of *fqgrep, seqkit grep, ripgrep, grep, awk* and *gawk*, using the *stat* command (Apple Inc. 2023b), and any difference investigated using the *diff* command (Apple Inc. 2023a). Furthermore, a custom utility, *spikeq* (Crosbie 2024b), was developed to generate synthetic FASTQ files with a known number of records and sequences with user-specified lengths that were spiked with a set of regular expressions a known number of times. This utility was used to test the performance of *grepq* and the aforementioned tools under controlled conditions.

Finally, a bash test script (see *examples/test*.*sh*, available at *grepq*’s Github repository) and a simple Rust CLI application, *predate* (Crosbie 2024a), were developed and utilised to automate system testing, and to monitor for performance regressions.

*grepq* has been tested on macOS 15.0.1 (Apple M1 Max) and Linux Ubuntu 20.04.6 LTS (AMD EPYC 7763 64-Core Processor). It may work on other platforms, but this has not been tested.

## Availability and documentation

*grepq* is open-source and available at *GitHub* (https://github.com/Rbfinch/grepq) and *Crates*.*io* (https://crates.io/crates/grepq).

Documentation and installation instructions for *grepq* are available at the same GitHub repository, and through the **-h** and **–help** command-line options, which includes a list of all available commands and options, and examples of how to use them. Example pattern files in plain text and JSON format are also provided, as well as test scripts. *grepq* is distributed under the MIT license.

## Conclusion

The performance of *grepq* was compared to that of *fqgrep, seqkit grep, ripgrep, grep, awk*, and *gawk* using the benchmarking tool *hyperfine*. For an uncompressed FASTQ file 874MB in size, containing 869,034 records, *grepq* was significantly faster than the other tools tested, with a speedup of 1797 times relative to *grep*, 864 times relative to *awk*, and 19 times relative to *ripgrep*. For a larger uncompressed FASTQ file (104GB in size, and containing 139,700,067 records), *grepq* was 4.4 times faster than *ripgrep* and marginally slower or of equivalent speed to *ripgrep* where the same large file was gzip-compressed. When coupled with its exceptional runtime performance, *grepq*’s feature set make it a powerful and flexible tool for filtering large FASTQ files.

## Acknowledgements

I’m grateful to my family for their patience and support during the development of *grepq*. I would also like to thank the developers of the *seq_io, regex, mimalloc* and *flate2* libraries for their excellent work, and the developers of the *hyperfine* benchmarking tool for making it easy to compare the performance of different tools. Finally, I would like to thank the authors of the *ripgrep* and *fqgrep* tools for providing inspiration for *grepq*.

